# High-throughput plasma proteomic platforms: Insights from a multi-ethnic Asian cohort

**DOI:** 10.1101/2025.10.24.682486

**Authors:** Akash Bahai, He Wang, Pritesh R Jain, Lucas D Ward, Oleksandr Boychenko, Zhihao Ding, Jun Frederiksen, Joanne Ngeow, Jimmy Lee, Elio Riboli, Jianjun Liu, Sebastian Maurer-Stroh, Enrico Petretto, Ching-Yu Cheng, Bernett Lee, Saumya Shekhar Jamuar, Nicolas Bertin, Claire Bellis, Max Lam, Weiling Zheng, Xueling Sim, Patrick Tan, John C Chambers, Nilanjana Sadhu

## Abstract

We compared three leading affinity-based and mass-spectrometry-based proteomic platforms (SomaScan11K, Olink Explore HT, Orbitrap Astral with Seer Proteograph [MS-Seer]) in a multi-ethnic Asian cohort, to inform biomarker discovery and functional genomic studies of global populations. We found limited overlap (1,740 proteins) across the three platforms, with modest correlations (0.34-0.10). SomaScan had lower missingness (<1%) and CV (<10%), compared to Olink (51%, 23%) and MS-Seer (12%, 19%). The new assays in Olink Explore HT (absent in Explore3072) primarily drove its higher missingness and CV. The number of phenotypic associations varied by trait, while the number of genetic associations (*cis-*pQTLs at P<5e-08) were similar for Olink and SomaScan. Protein levels differed between ethnicities, with SomaScan identifying more ethnicity-differentiated proteins than Olink (FDR<0.05). Finally, SomaScan ANML normalization attenuated biologically relevant associations in our study. These findings underscore the importance of platform evaluation and data normalization strategies for application in large-scale, diverse population cohorts.

## Introduction

Population-scale proteomics has heralded a new era in translational medicine, facilitating functional validation of genomic discoveries and gene target prioritization^1^. As a molecular phenotype, circulating proteins capture both genetic and environmental influences, providing insights into healthy and disease-perturbed pathways^2^. Large scale proteomic studies carried out in the UK Biobank^2^, DeCODE^3^, ARIC^4,5^, and INTERVAL^6^ studies have capitalized on proteomic profiling to identify robust biomarkers for incident disease, risk stratification and disease progression, and to identify potential therapeutic targets in novel pathways. Whilst these studies have furthered the pursuit of precision medicine^7^, they are also notable for their focus on European and North American populations. There is a widely recognized need for comparable, well-powered studies in other global population groups, both as an opportunity to understand shared and population specific disease pathways, and to evaluate the generalizability of proteomic biomarkers across populations.

Mass spectrometry (MS) and multiplexed affinity-based assay methods have emerged as the two leading, complementary strategies for quantification of proteomic variation in samples from large-scale population-based studies. While MS has been considered the gold-standard for highly specific quantification of peptides, proteins, and proteoforms, many MS-based methods are recognized to have limited sensitivity, dynamic range and throughput^8,9^. State-of-the-art MS technologies such as the Orbitrap Astral (Thermo Fisher Scientific) and the timsTOF Pro (Bruker), have substantially improved assay performance, and may enable quantification of many thousands of plasma proteins from a single sample, especially when combined with innovative approaches to protein enrichment and depletion such as the Seer Proteograph or PreOmics ENRICH-iST^9^. Although preliminary results are encouraging^10,11^, there is currently limited experience of these advanced MS-based approaches, and their suitability for use in large scale population studies remains to be determined.

In parallel, the aptamer-based SomaScan and antibody-based Olink assays have continuously improved their respective technologies, including increased depth and throughput. The most recent iterations of the SomaScan and Olink platforms can now assay 9,655 and 5,416 unique human proteins respectively, in a high-throughput and cost-effective manner. However, these ligand binding assays are recognized to have systematic differences in protein coverage, as well as specificity and reproducibility of their assays^12^. These differences in performance may contribute to limited cross-platform reproducibility, and impact biological insights. Furthermore, the performance of the newly added content in the latest generation of these affinity-based assays, has had limited technical or biological evaluation^11,13,14^.

To advance understanding of disease pathways in the genetically, environmentally and culturally distinct populations of Asia, we have established a comprehensive longitudinal study (PRECISE-SG100K) comprising over 100,000 men and women living in South-East Asia. In preparation for planned proteomic profiling of all 100,000 participants, we carried out a pilot study, to evaluate the performance of the latest generation of MS and affinity-based assays.

## Results

We performed a comparative analysis of the aptamer-based SomaScan 11K platform (SomaScan11K), the antibody-based Olink Explore HT platform (OlinkHT), and the Seer Proteograph XT workflow for nanoparticle-based enrichment coupled with data-independent acquisition by the Orbitrap Astral mass spectrometer (MS-Seer). The primary objective of our study was to evaluate their technical reproducibility, precision, specificity, and discovery potential for future large-scale application in our large multi-ethnic cohort, the PRECISE-SG100K study.

### Study cohort characteristics

Plasma proteomic profiles were generated in 800 samples from 500 participants, adopting a design that included analysis of both technical as well as longitudinal replicates (**Figure 1**). All 800 samples were assayed using SomaScan11K, of which 516 samples from 216 participants were assayed using OlinkHT, and 100 samples from 50 participants were assayed using MS-Seer. Sample size varied because of limitations in resource availability, however, inclusion of overlapping samples provides opportunity for valuable orthogonal benchmarking. All MS-Seer samples were also analyzed by both SomaScan11K and OlinkHT. The study cohort included people of Chinese (East Asian), Indian (South Asian), and Malay (South-East Asian) background, with an average age of 51 ± 12 years, and 54% females. Further details of participant characteristics are shown in **Supplementary Table 1**.

**Figure 1.**
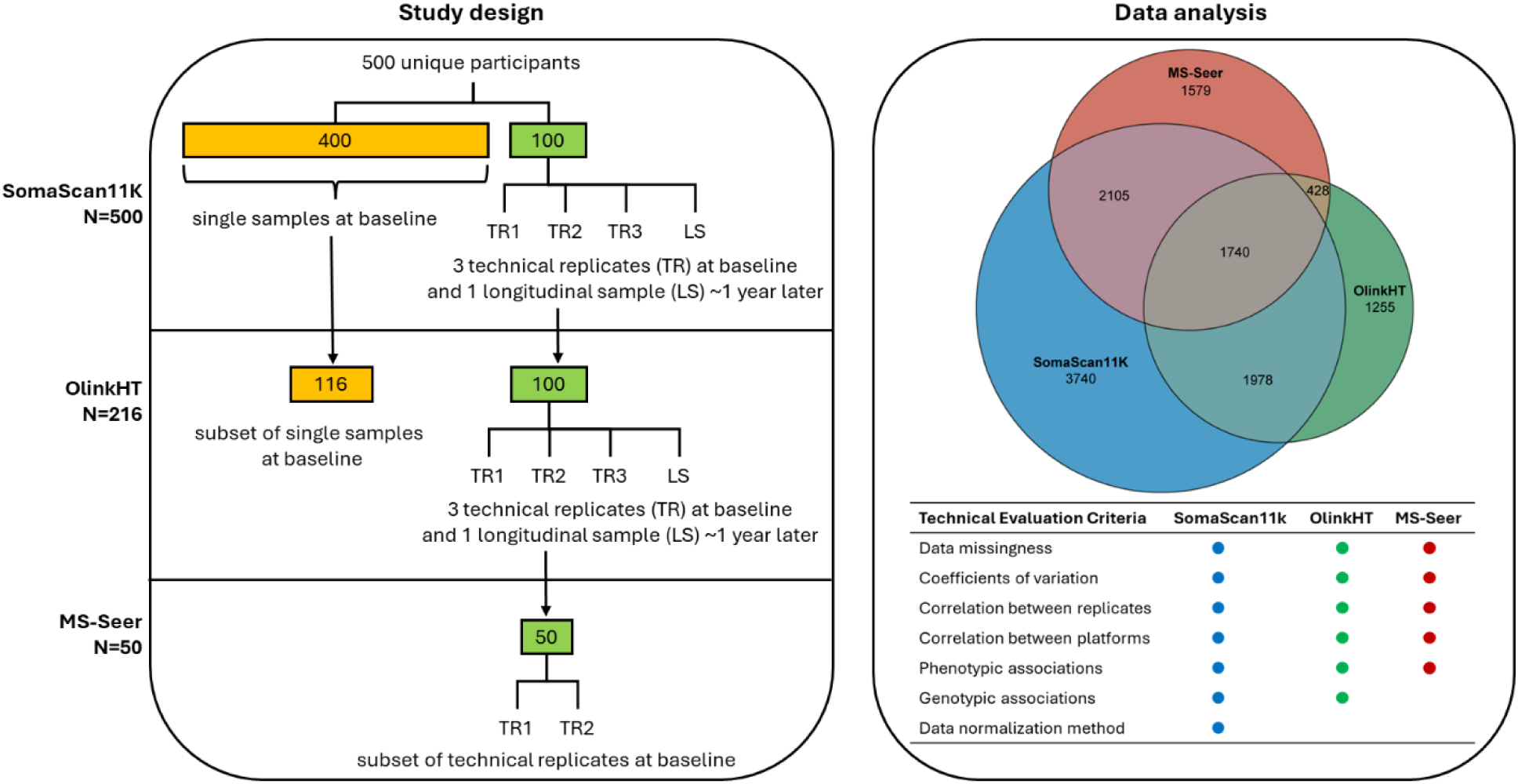
Study design for comparison of plasma proteomic platforms. (Left) Hierarchical sampling workflow of the cohort for the SomaScan11K, OlinkHT, and MS-Seer platforms. (Right, top) Venn diagram depicting the number of proteins uniquely and commonly quantified by three platforms. (Right, bottom) Summary of the seven technical evaluation criteria applied in this study.

### Proteomic profiling by the three platforms

SomaScan11K returned results for 10,675 SOMAmers (mapping to 9,563 unique proteins as identified by UniProt IDs), while OlinkHT returned 5,405 features (5,401 UniProt IDs) and MS-Seer identified 5,945 protein groups (5,852 UniProt IDs). Overall, 12,825 unique proteins were identified by one or more of the three platforms (**Figure 1**). There were 3,718 proteins shared between SomaScan11K and OlinkHT, 3,845 shared between SomaScan11K and MS-Seer, and 2,168 shared between OlinkHT and MS-Seer. There were only 1,740 proteins common across all three platforms (**Supplementary Table 2**).

Data generated by each of the three platforms effectively distinguished between experimental and control samples (**Extended Data** Figure 1a-g). There were no obvious batch or plate effects (**Extended Data** Figure 1h-n). There were no outlier samples in the OlinkHT and MS-Seer data, and two outliers identified in the SomaScan11K data had not been assayed on OlinkHT or MS-Seer (**Extended Data** Figure 2). For OlinkHT and MS-Seer, the analyses were based on plate-scaled data. SomaLogic additionally apply a proprietary method called Adaptive Normalization by Maximum Likelihood (ANML) to their plate-scaled data, which normalizes measurements to a healthy U.S. population reference by iteratively estimating dilution-bin-specific scale factors from protein expression values within two standard deviations of the reference. The resulting scale factors are then applied to their respective SOMAmer reagents, to enhance technical performance and promote run-to-run consistency^15,16^. For SomaScan11K, we therefore had two sets of data: (i) pre-ANML, which is plate-scaled data; and (ii) ANML, which is the final normalized dataset recommended by SomaLogic that applies ANML in addition to plate-scaling. We evaluated the performance of both ANML and pre-ANML data for SomaScan11K.

### Technical performance

In the SomaScan11K (500 samples and 10,675 aptamers) dataset, the median proportions of proteins with measurements below the estimated limit of detection (LoD) per sample were 0.7% (pre-ANML) and 2.6% (ANML). In contrast, a median of 11.8% of proteins for MS-Seer (46 samples and 5,945 proteins) and 51.4% of proteins for OlinkHT (216 samples and 5,405 assays) had measurements below LoD (**Figure 2a; Supplementary Figure 1**). At a threshold of ≤20% missingness to define successful protein quantification, we found that 96% and 80% of proteins were successfully called in the SomaScan11K and MS-Seer datasets. In contrast, only 40% of Olink assays passed this permissive calling threshold (**Figure 2b**).

**Figure 2.**
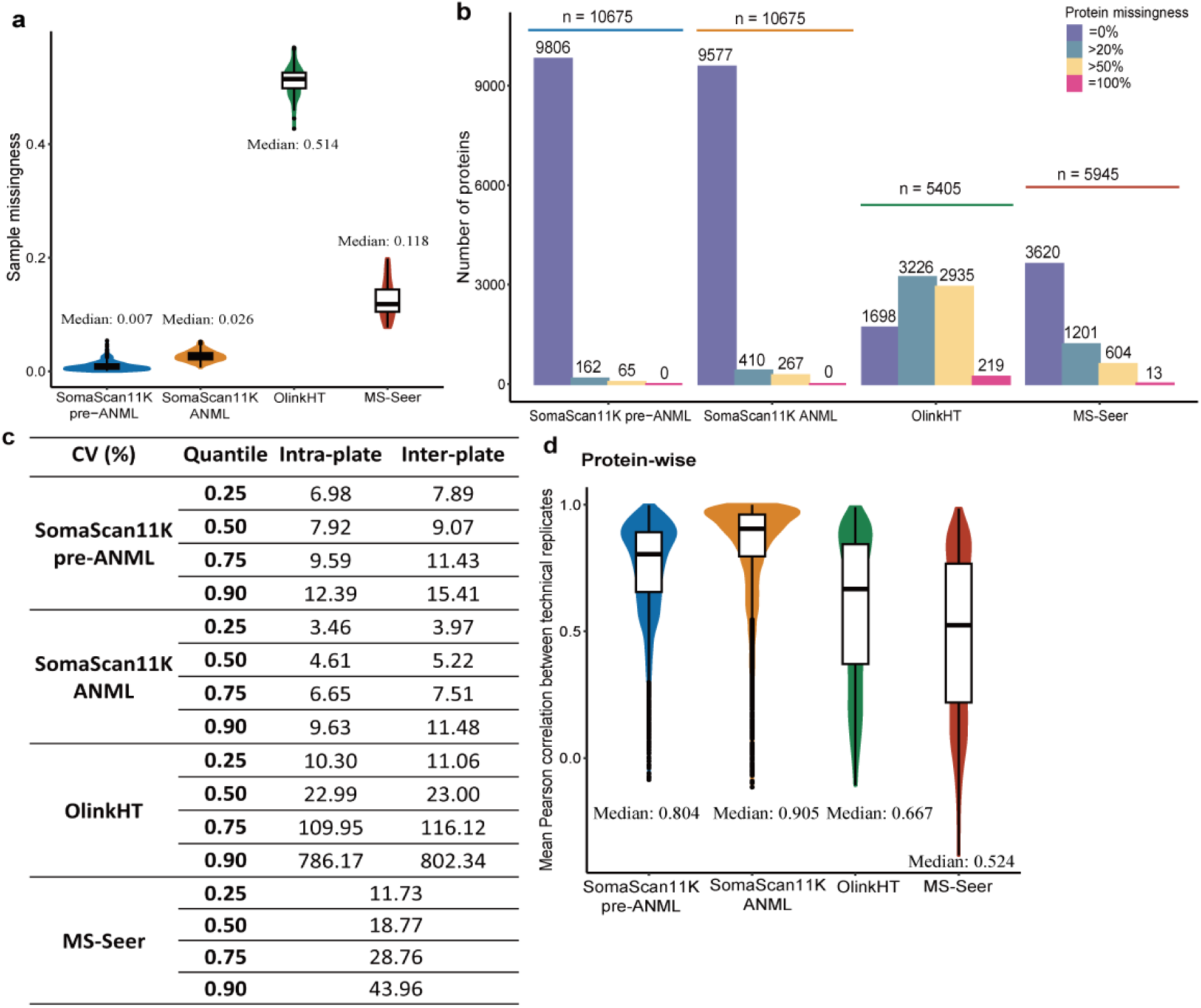
Technical evaluation across SomaScan11K, OlinkHT, and MS-Seer proteomics platforms. (a) Sample-level measurability: violin plots depicting sample missingness for three platforms. (b) Protein-level measurability: bar graph showing the numbers of proteins with different levels of missingness across the platforms, categorized by different thresholds. (c) Precision of measurements: coefficients of variation for SomaScan11K, OlinkHT, and MS-Seer platforms. (d) Technical reproducibility: violin plots showing the mean protein-wise Pearson correlation between technical replicates for three platforms.

Coefficients of variations (CVs) between technical replicates showed that median intra-and inter-plate CVs were lowest for the SomaScan11K assay, for both pre-ANML and ANML data (pre-ANML: intra 7.9%, inter 9.1%; ANML: intra 4.6%, inter 5.2%; **Figure 2c**). In contrast, CVs for technical replicates were higher for both OlinkHT (intra: 23.0%, inter: 23.0%) and MS-Seer (overall: 18.8%). CVs for longitudinal samples collected 1 year apart were also higher for OlinkHT (28.3%) compared to SomaScan11K (pre-ANML: 10.4%, ANML: 6.3%; **Supplementary Table 3**). Similarly, both SomaScan11K pre-ANML and ANML exhibited higher correlation between technical replicates, compared to OlinkHT and MS-Seer (**Figure 2d; Supplementary** Figure 2). Median pair-wise correlation for protein concentrations across technical replicates was 0.80, 0.91, 0.67, and 0.52, for SomaScan11K pre-ANML, ANML, OlinkHT, and MS-Seer, respectively (**Figure 2d**). Overall findings remained unchanged when restricting these analyses to common samples (45 pairs) and common proteins (1,740) across the three platforms (**Supplementary Figure 3**).

### Orthogonal confirmation

Next, we calculated pairwise Spearman correlations for common proteins shared between platforms (SomaScan11K and MS-Seer [3,845 proteins, 46 samples], OlinkHT and MS-Seer [2,168 proteins, 46 samples], and SomaScan11K and OlinkHT [3,718 proteins, 216 samples], **Supplementary Table 4**). Median values for the correlation between the affinity-based assays and MS-Seer assay were relatively low (SomaScan11K pre-ANML: 0.27, SomaScan11K ANML: 0.26, OlinkHT 0.34; **Figures 3a-c; Supplementary Table 5**). More than half of the common proteins in MS-Seer-SomaScan11K ANML (53.4%) and MS-Seer-OlinkHT (56.2%) comparisons had higher correlations than the 99th-percentile of their null distributions, whereas the proportion was lower for MS-Seer-SomaScan11K pre-ANML (38.1%).

**Figure 3.**
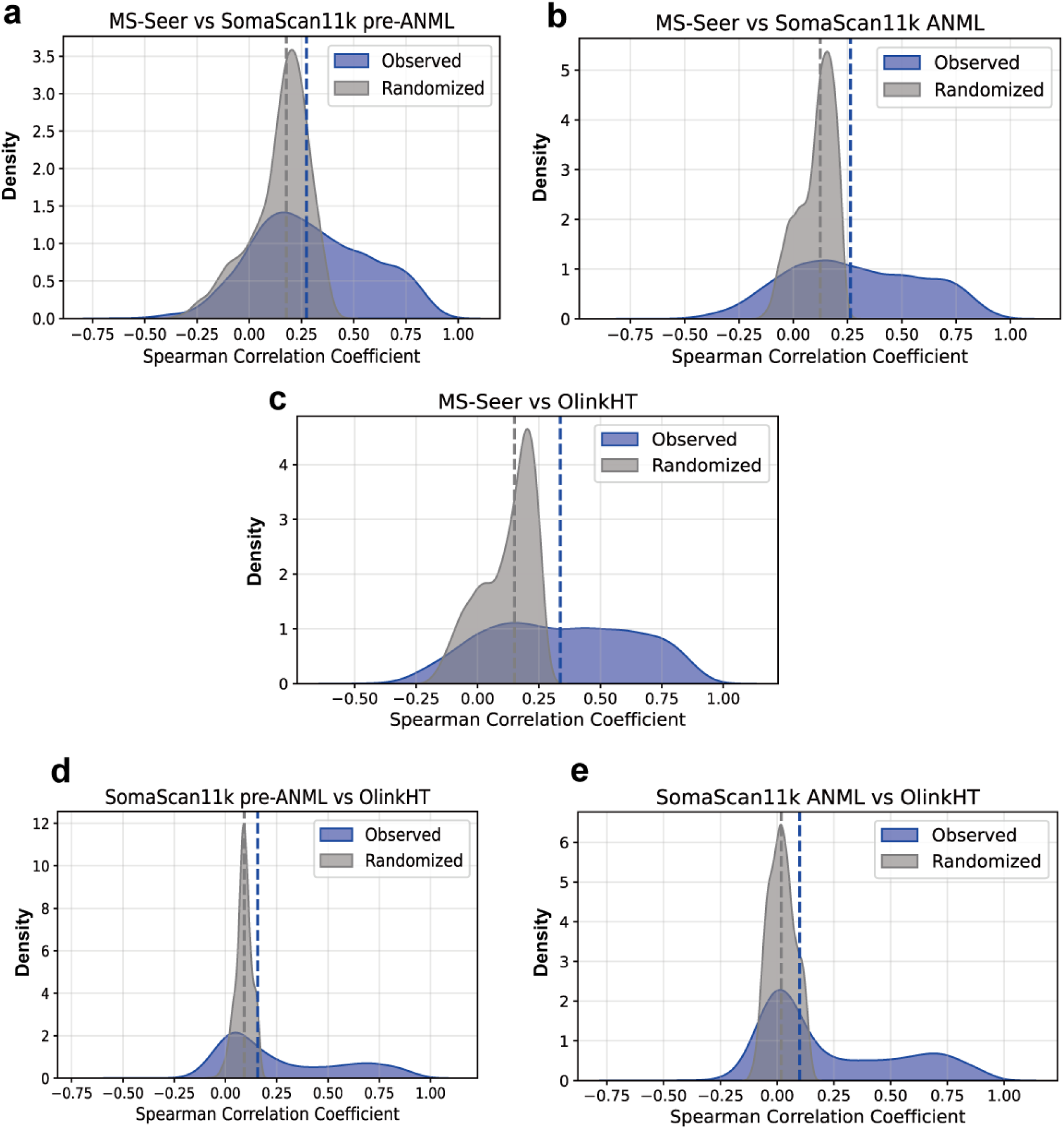
The distribution of protein-wise Spearman correlation between different platform pairs. The distribution in blue is the observed correlation and the grey distribution is the randomized (background) correlation. The blue and grey dotted lines represent the median of each distribution, respectively. (a) Correlation distribution of shared proteins between MS-Seer and SomaScan11K pre-ANML data (b) Correlation distribution of shared proteins between MS-Seer and SomaScan11K ANML data (c) Correlation distribution of shared proteins between MS-Seer and OlinkHT data (d) Correlation distribution of shared proteins between SomaScan11K pre-ANML and OlinkHT data (e) Correlation distribution of shared proteins between SomaScan11K ANML and OlinkHT data.

The median correlations between SomaScan11K and OlinkHT were also low (pre-ANML: 0.16, ANML: 0.10; **Figures 3d,e; Supplementary Table 5**). There was a bimodal distribution in the correlations with the largest peak at ∼0.0 and a secondary peak at ∼0.75, for both ANML and pre-ANML data **(Figures 3d,e)**. For SomaScan11K-OlinkHT correlations, 44.7% of ANML and 48.5% of pre-ANML proteins exceeded the 99th-percentile threshold of the null distribution, indicating that approximately half of the overlapping proteins exhibit high cross-platform concordance.

### Phenotypic and genotypic associations

Our analyses focused on the 216 samples with data for both SomaScan11K and OlinkHT using either (1) all proteins assayed or (2) the common set of proteins assayed by both platforms. We found that SomaScan11K pre-ANML data had the highest number of proteins associated with sex and BMI, while OlinkHT had the highest number of associations with age (**Table 1a)**. For all three phenotypes, we noted marked differences in the number of associations identified between SomaScan11K pre-ANML and ANML data, especially for BMI (pre-ANML: 4,361, ANML: 1,234 associations at FDR<0.05; **Table 1a; Supplementary Tables 6-7)**.

**Table 1.** Comparison of phenotypic and genotypic associations for SomaScan11K pre-ANML, SomaScan11K ANML and OlinkHT. (a) The total number of associations (FDR < 0.05) with Age, Sex, and BMI. (b) The total number of *cis-*pQTLs (p<5e-08), and *trans-*pQTLs (p<5e-08/number of UniProt IDs). The numbers of associations are based on number of unique UniProt IDs with at least one significant assay association. The numbers of assay associations are displayed in brackets.

| a) |  | # UniProt ID | # Assay | Age | Sex | BMI |
| --- | --- | --- | --- | --- | --- | --- |
| All proteins,<br>Common<br>samples | SomaScan11K<br>pre-ANML | 9563 | 10675 | 445 (486) | 1882 (1986) | 4361 (4604) |
|  | SomaScan11K<br>ANML | 9563 | 10675 | 652 (697) | 495 (528) | 1234 (1304) |
|  | OlinkHT | 5401 | 5405 | 704 | 401 | 1145 (1146) |
| Common<br>proteins,<br>Common<br>samples | SomaScan11K<br>pre-ANML | 3718 | 4424 | 284 (320) | 913 (990) | 1621 (1781) |
|  | SomaScan11K<br>ANML | 3718 | 4424 | 365 (401) | 328 (356) | 580 (638) |
|  | OlinkHT | 3718 | 3722 | 629 | 391 | 1008 (1009) |
| b) |  | # UniProt ID | # Assay | *cis*-pQTLs | *trans*-pQTLs |  |
| All proteins,<br>Common<br>samples | SomaScan11K<br>pre-ANML | 9563 | 10675 | 298 (349) | 127 (129) |  |
|  | SomaScan11K<br>ANML | 9563 | 10675 | 351 (416) | 191 (193) |  |
|  | OlinkHT | 5401 | 5405 | 300 (305) | 16 (17) |  |
| Common<br>proteins,<br>Common<br>samples | SomaScan11K<br>pre-ANML | 3718 | 4424 | 186 (219) | 37 (38) |  |
|  | SomaScan11K<br>ANML | 3718 | 4424 | 218 (262) | 59 (60) |  |
|  | OlinkHT | 3718 | 3722 | 261 (265) | 15 (16) |  |

Cross-platform comparison of phenotypic associations including MS-Seer was limited in power owing to the small number of common samples (n=46), with no detected associations with sex. Known markers of sex, including leptin, sex-hormone binding globule, pregnancy zone protein, were nominally associated across all three platforms at P<0.05. For age, and BMI, MS-Seer had the lowest number of associations (age=11, BMI=50) compared to SomaScan11K (pre-ANML: age=17, BMI=237; ANML: age=23, BMI=82) and OlinkHT (age=71, BMI=60) at FDR<0.05 (**Supplementary Tables 8-10**).

We also performed genome-wide association analysis for all analytes in SomaScan11K and OlinkHT in the common set of 216 samples. Since SomaScan11K has multiple aptamers targeting the same protein, we compared the number of unique proteins with at least one significant assay association, to avoid overestimation. We found that SomaScan11K ANML identified the highest number of proteins with at least one cis-pQTL (pre-ANML: 298, ANML: 351, OlinkHT: 300 at p<5e-08; **Table 1b**). Approximately 45% of the cis-pQTLs identified by each platform are also reported as cis-eQTLs (p<0.05) in the eQTLGen dataset^17^. Fine-mapping revealed a greater number of unique causal variants for SomaScan11K compared to OlinkHT (pre-ANML: 123, ANML 142, OlinkHT: 117; **Supplementary Table 11)**. Finally, SomaScan11K (both pre-ANML and ANML) identified a higher number of proteins with trans-pQTLs (at p<5e-08/number of UniProt IDs) compared to OlinkHT (**Table 1b**).

### Proteomic profiles vary between Asian ethnic groups

Global PCA of all analytes across all samples for OlinkHT (5,405 analytes, 216 samples) and SomaScan11K (pre-ANML and ANML, 10,675 analytes, 500 samples) data revealed that the Indian and Malay proteomes were significantly different from that of the Chinese (Hotelling’s T2 test P<0.05; **Supplementary Table 12, Supplementary Figure 4**). Differences between the Indian and Malay groups were also evident in the SomaScan11K dataset (N=500, **Supplementary Table 12**). SomaScan11K identified a greater number of ethnicity-associated proteins compared to OlinkHT, even in the common set of 216 samples (**Supplementary Table 13**). There were 191 proteins robustly identified as ethnicity-differentiated across all three datasets (FDR <0.05 for at least one pairwise comparison, and same direction of effect for the corresponding comparison across OlinkHT, and SomaScan11K Pre-ANML and ANML data, **Supplementary Table 14**). Our analysis captured proteins previously reported to vary across ancestries, such as Leptin, which was found to be highest in Indians and lowest in Chinese, reflecting known differences in BMI and cardiovascular outcomes between these ethnic groups^18,19^. It also identified ethnic differences in Lipopolysaccharide-binding protein (LBP), a glycoprotein involved in immune response and atherogenesis^20,21^. LBP levels were found to be higher in Indian and Malay individuals compared to the Chinese, and strongly positively correlated with cardiovascular disease risk factors (BMI, C-reactive protein, triglyceride, and total cholesterol to high density lipoprotein ratio levels; **Supplementary Figure 5**).

### Stratification by assay content

The SomaScan and Olink affinity-based platforms have both evolved in a stepwise manner to increase the number of proteins assayed. To understand how this might relate to assay performance, we compared results for SomaScan and Olink data, stratified by assay release (SomaScan sets 1 to 3: 5K assays, 7K minus 5K assays, 11K minus 7K assays; Olink sets 1 to 3: Explore 1536 assays, Explore 3072 minus Explore 1536 assays, Explore HT minus Explore 3072 assays; see **Methods**). We found that missingness and CV were consistently low, and that technical reproducibility was consistently high, across assay sets for SomaScan **(Supplementary Figure 6).** The number of phenotypic associations from each set of SomaScan was also proportional to the total number of proteins in each set, but a majority of the genotypic associations were from set 1 (**Supplementary Tables 15-16**). In contrast, we noted that missingness and CVs were progressively higher, and reproducibility progressively lower, for the newer Olink assay content. For example, missingness increased from 14.4% in set 1 to 76.7% in set 3, while CV increased from ∼9% in set 1 to ∼40% in set 3 **(Supplementary Table 17).** We also noted that there were fewer phenotypic and genotypic associations in the most recent Olink assay content. Irrespective of assay version, above-LoD proteins and higher-abundance proteins had better technical performance in both SomaScan and Olink platforms (**Supplementary Figures 7-9**).

### Comparison between SomaScan11K pre-ANML and ANML data

From our technical and biological evaluations, we observed that the SomaScan11K ANML dataset generally has a lower CV, higher correlation between technical replicates, a higher number of age-associated proteins, and more pQTLs. However, the ANML data also show higher missingness, weaker orthogonal confirmation, and fewer associations with sex and BMI. We confirmed this through recalculation of genotypic and phenotypic associations using all 500 samples and 10,675 aptamers from the SomaScan11K platform for ANML and pre-ANML data. Even with this larger dataset, ANML retained a higher number of pQTLs and associations with age, whereas pre-ANML demonstrated significantly more associations with sex and BMI (**Supplementary Table 18**).

To evaluate the impact of ANML on data content, we first estimated the pair-wise Spearman correlation between our pre-ANML and ANML data. We found that 52.6% of proteins had a correlation below 0.8 (**Figure 4a**) between their pre-and post-ANML values, indicative of changes in the data structure owing to ANML scaling. We also observed that the ANML scaling factors strongly correlated with a range of phenotypes in our dataset, especially BMI, common cardiometabolic markers, and self-reported ethnicity (**Figure 4b, Extended Data** Figure 3), raising the possibility that ANML scaling factors are partly determined by physiological variation between participants. In keeping with this, we find that adiposity and related cardiometabolic phenotypes (e.g., CRP, BMI and insulin) were most predictive of ANML scaling factors when ranked based on their random forest-based Boruta feature selection method (**Extended Data** Figure 4).

**Figure 4.**
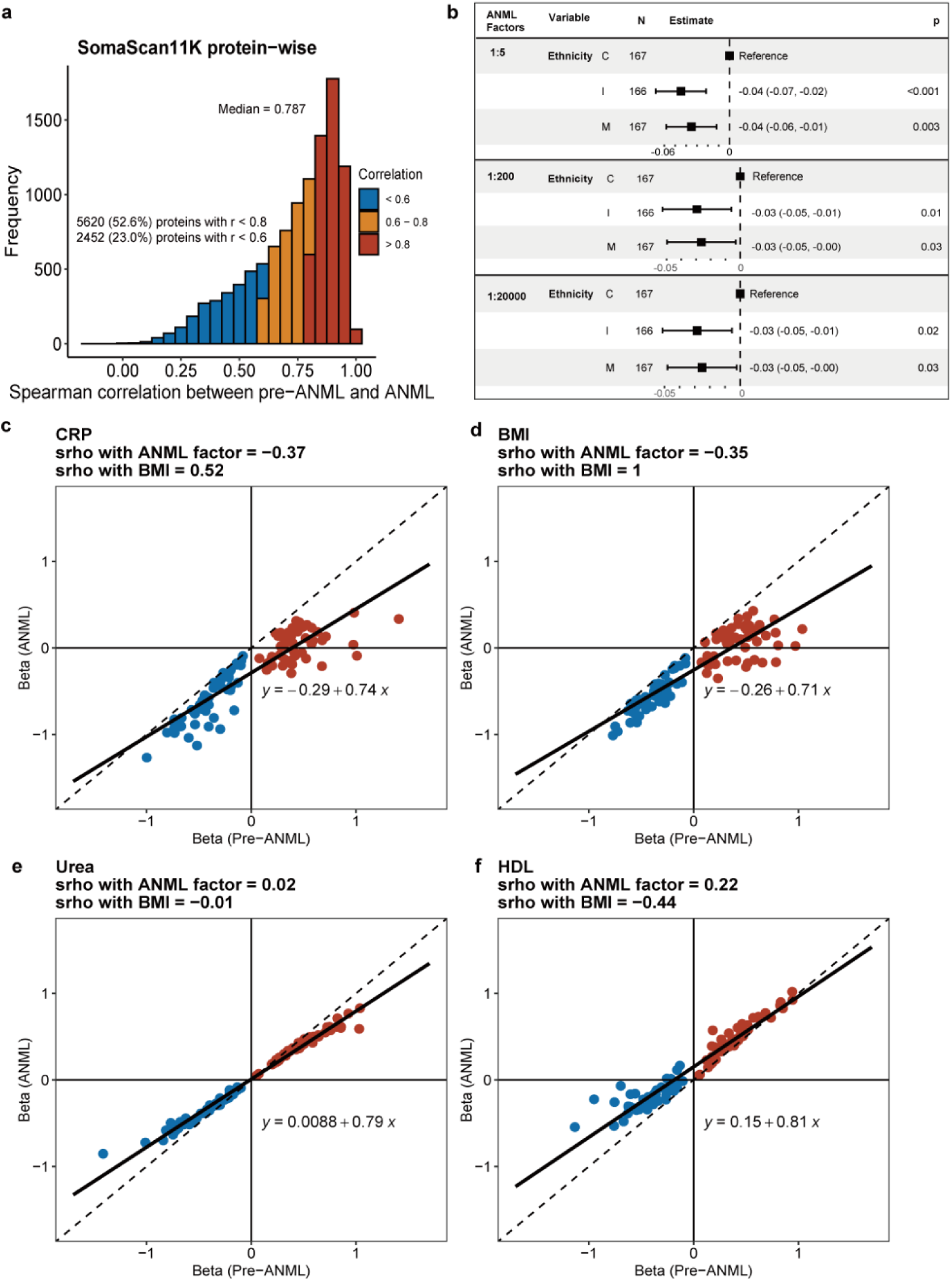
Comparison of SomaScan11K pre-ANML and ANML data. (a) Histogram showing the distribution of protein-wise Spearman correlations between pre-ANML and post-ANML SomaScan11K data. (b) The forests plots depicting the results from fitted linear models (ANML scaling factors ∼ ethnicities). (c-f) Effect size comparison for 4 representative phenotypes between pre-ANML and ANML of synthetic proteins generated by simulation method 1: (c) CRP, (d) BMI, (e) Urea and (f) HDL. The Spearman correlations between each phenotype and ANML scaling factor/BMI are shown in each panel title. The dotted and solid lines represent y = x and fitted line, respectively. Red: pre-defined positive effect size; Blue: pre-defined negative effect size.

### Analysis of SomaLogic ANML normalization through data simulation

To investigate the impact of ANML scaling on associations between protein levels and phenotype, we created synthetic relationships through spike-ins of the pre-ANML data (see **Methods**). The synthetic relationships were designed to mimic associations with 20 different phenotypes, across a plausible range of epidemiological effects. The 20 phenotypes were selected to include phenotypes negatively, minimally or positively associated with ANML scaling factors (e.g. BMI, urea, and HDL, respectively). We then compared the results for the associations of the spiked proteins with the chosen phenotypes between the pre-ANML and ANML datasets.

We found that ANML led to a systematic shift in both the effect size and significance level of the association between protein level and phenotype (**Figures 4c-f; Extended Data Figures 5-6; Supplementary Figures 10-11**). In keeping with our initial hypothesis, the impact of ANML on the association statistics appeared to mirror the magnitude and direction of the correlation between the phenotypes and ANML scaling factors. For phenotypes that were highly negatively correlated with ANML scaling factors (e.g. CRP, BMI), beta coefficients were consistently lower (more negative) in the ANML analyses, to an extent that even reversed the direction of effect. Conversely, phenotypes that were highly positively correlated with ANML scaling factors (e.g. HDL) became higher (more positive). Phenotypes that showed only weak correlations with ANML scaling factors (e.g. urea) were minimally affected by ANML normalization.

## Discussion

We evaluated three proteomic platforms designed to deliver deep proteomic profiling suitable for population-scale studies. There were ∼13,000 proteins quantified across the three platforms, however the overlap was only 1,740 proteins. As an initial key observation, we note that each platform measures a different fraction of the plasma proteome, indicating complementary information content for population studies.

Comparing technical performances, SomaScan11K had the lowest missingness, lowest CVs, and highest correlation between replicates, indicating stable and replicable measurements. OlinkHT’s high missingness and high CVs were found to be primarily driven by the newest set of assays in their latest generation of the Explore panel. This observation is in agreement with another recent study that found the missingness and CV for the Olink Explore HT panel to be almost twice that of the Olink Explore 3072 panel^13^. These findings may be explained by the fact that the newer assays in Olink are largely representative of non-secretory proteins that might only present at very low levels in circulation unless in a specific disease state, leading to poor discoverability in generally healthy plasma and large variability in measurements using Olink’s current method of quantification. MS-Seer, on the other hand, had acceptable data missingness, and modest precision reflected by its higher CVs and lower protein-wise correlation between replicates in comparison with SomaScan11K.

Pair-wise correlation for the limited set of shared proteins, revealed that OlinkHT had a higher median correlation with MS-Seer compared to SomaScan11K. We further observe that the correlation between OlinkHT and MS-Seer is significantly higher in the subset of older assays (equivalent to Olink Explore 1536), as well as in the above LoD subset of analytes. Correlation between Olink and SomaScan platforms have been widely discussed^13–15,22–24^. We compare data from the latest generations of these two platforms and find a median correlation of 0.16 (pre-ANML) and 0.10 (ANML), lower than previously reported^13,14^. However, the distribution of the correlation remained similar to previous reports consisting of a bimodal curve with two peaks – one at 0.0, and the other at 0.75. When restricting the analysis to data above LoD for each platform, we find that the correlation between Olink and SomaScan increases to 0.46 (pre-ANML) and 0.43 (ANML).

We evaluated phenotypic and genotypic associations with common phenotypes such as age, sex, BMI. We find that above LoD analytes contribute to a majority of the associations for both OlinkHT and SomaScan11K. Overall, OlinkHT has a greater number of age-specific markers while SomaScan11K yields more sex-and BMI-specific markers. This may reflect differences in the composition of each platform’s marker panel, with each capturing a distinct subset of the proteome. We observe a similar number of proteins with *cis-*pQTL signals across the two platforms, however, *trans-*pQTL discovery was higher in SomaScan11K compared to OlinkHT. The number of *trans-*pQTLs discovered for SomaScan is relatively high given the limited sample size of 216^2^, raining the possibility of protein-protein interactions or off-target effects. Further studies are needed to better understand the nature of the *trans-*pQTLs.

We noted substantial differences in the distribution of protein levels and association test results, between the SomaScan11K pre-ANML and ANML data, in agreement with previous reports^15,23,24^.

The ANML method aims to adjust for unwanted technical variations and reduce batch effects in large-scale proteomic studies - the reference population for which may not reflect the ethnic diversity of the SG100K cohort^25^. We find that for 52.6% of the proteins, pre-ANML and ANML concentrations are only weakly correlated. We show that ANML scaling factors are correlated with ethnicity, adiposity and multiple cardiometabolic phenotypes, and that ANML normalization systematically shifts data distributions, with the correlation between the phenotype and the ANML scaling factor predicting the extent and direction of this effect on phenotypic associations. Additionally, pre-ANML data demonstrates higher orthogonal confirmation in our data, consistent with prior studies reporting higher correlation between pre-ANML and Olink data compared to ANML data^15,23^. Our evaluation of the effect of ANML normalization on biological discovery extends on these observations, providing valuable insights and suggesting that while ANML normalization may improve technical metrics, it may also be removing biological signals from datasets.

Our comparative analysis finds that the SomaScan11K assay (both pre-ANML and ANML) has lower missingness and higher reproducibility than OlinkHT and MS-Seer assays - two critical measures of technical performance. We observe that many of the new OlinkHT assays have poorer technical performance than the assays in Olink’s earlier generation Explore panels. While OlinkHT and SomaScan11K exhibit distinct strengths in biological discovery, we identify more proteins with ethnicity-associated differences using SomaScan11K than OlinkHT. Additionally, we demonstrate that ANML normalization, a proprietary algorithm applied by SomaLogic as the final step in their data generation, may be removing biologically relevant signals. Nevertheless, we note that the three platforms continue to have largely non-overlapping content, indicating that population-scale studies are likely to benefit from the implementation of a combination of technologies to maximize discovery potential. Our findings will enable rationale design and platform selection for proteomic studies focussed on advancing identification of biomarkers and disease pathways that are relevant to global populations.

## Methods

### Study design

Baseline samples from 216 participants were randomly selected from the PRECISE-SG100K cohort (https://www.npm.sg/partners/precise-sg100k/) to be run on two different platforms: Olink Explore HT and SomaScan 11K Assay. For 100 of these samples, three technical replicates were included in the study, as well as a longitudinal sample collected during a follow-up visit, approximately a year later. An additional 284 baseline samples selected randomly from the cohort were run on the SomaScan platform. Of the 100 baseline samples with replicates, a random subset of 50 baseline samples in duplicates were run on the Thermo Fisher Orbitrap Astral Mass Spectrometer (MS) instrument coupled with Seer Proteograph XT workflow. Thus, a total of 516 samples were run using Olink Explore HT, 800 samples using SomaScan 11K, 100 samples using Orbitrap Astral MS.

All assays were carried out in EDTA plasma separated from venous whole blood, collected from participants in the fasting state. Blood samples were collected by a certified phlebotomist, during the morning of the participant’s visit. Samples were centrifuged within four hours and plasma was stored at-80 degrees Celsius before analysis. The overall study samples were selected such that they were age-, sex-matched across the three ethnic groups, i.e., Chinese, Indian and Malay. Samples were randomized across plates. For samples with replicates, two technical replicates were on the same plate to enable assessment of intra-plate CVs, and the third technical replicate was on a different plate to enable inter-plate CVs. Longitudinal replicates were on the same plate as the third technical replicate to enable longitudinal CVs. It was also ensured that replicates were not adjacent to each other.

### Aptamer-based proteomics platform

800 samples from 500 participants from PRECISE-SG100K, along with 32 buffer samples (without added sample), 50 calibrators, and 30 QC samples (provided by SomaLogic from pooled healthy donor controls), divided into 10 plates from 2 batches, were analyzed using the SomaScan 11K Assay v5.0^26^ (*SomaLogic, Inc.*) at the SomaLogic Boulder Lab. This assay comprises 10,776 SOMAmers, which mapped to 9,655 UniProt IDs provided by SomaLogic; the remaining SOMAmers are controls, including hybridization control elution SOMAmers used in normalization and others targeting non-human proteins. To cover a broad range of protein concentrations, SOMAmers are grouped into dilution bins of 1:5, 1:200, and 1:20,000. Each sample required 130 μL of human plasma. Data processing, normalization, and quality control were conducted by SomaLogic, following sequential procedures including hybridization normalization, plate scaling and calibration^27^. SomaLogic also applied additional adaptive normalization by maximum likelihood (ANML) to the QC and experimental samples. To understand how ANML influences SomaScan11K data, we included both pre-ANML and ANML data in our comparison. One buffer sample failed QC evaluation and was excluded; all experimental samples passed QC. For all analysis, we used log2-transformed relative fluorescence units (RFU) for SomaScan11K data.

### Antibody-based proteomics platform

Antibody-based proteomic profiling was performed at the Olink Singapore Lab using the Olink Explore HT platform^28^ (*Olink Proteomics*). This assay employs proximity extension assay technology to measure 5,420 analytes (mapped to 5,416 UniProt IDs provided by Olink) using only 2 μL of plasma. To encompass a broad range of protein abundances, assays were grouped into dilution series of 1:1, 1:10, 1:100, 1:1,000, and 1:100,000. A total of 516 samples from 216 participants from PRECISE-SG100K were analyzed. The 60 external control samples (including 12 negative controls, 30 plate controls, and 18 sample controls from Olink), along with experimental samples, were distributed across 6 plates in 3 batches. Three types of internal controls—incubation control, extension control, and amplification control—were added to all samples. Two samples were excluded due to QC failure. Normalized Protein eXpression (NPX) values were then generated in two main steps: first, sequencing counts of each assay in a sample were divided by the extension control count for that sample and log2-transformed; second, intensity normalization was performed by subtracting the median of the experimental samples within each plate for each assay.

### Mass spectrometry-based proteomics platform

MS-based untargeted proteomic profiling using the Seer Proteograph XT workflow^10^ (*Seer, Inc.*) coupled with the Orbitrap Astral MS instrument^29^ (*Thermo Fisher Scientific*) was performed at the Seer Redwood Lab. All samples—including 100 plasma samples from 50 participants from PRECISE-SG100K and 15 pooled plasma replicates from Seer - were processed (each sample was diluted from 120 μL to a total input volume of 240 μL for processing) on a Seer SP100 automation instrument across three Proteograph XT Assay plates; five samples were lost due to trypsin dispensing errors. Peptides were then separated using a Thermo Scientific Vanquish Neo UHPLC system under nanoflow conditions at 1,000 nL/min with a 21-minute gradient. Mass spectrometry analysis was conducted in data-independent acquisition mode on the Orbitrap Astral instrument, using a 400-ng injection amount and a total acquisition time of 26 minutes per run. Each sample was run twice within each plate: once with XT Nanoparticle Suspension A and once with Suspension B.

### Mass spectrometry data processing

MS raw data were processed using DIA-NN^30^ (version 1.8.1) with a library-free search based on the Homo sapiens UniProt reference database (accessed December 2022). Peptide and protein intensities were quantified in match-between-runs mode using the following flags: `--mass-acc-ms1 3, --mass-acc 8, --qvalue 0.01, --matrices, --missed-cleavages 1, --met-excision,--cut K*,R*, --smart-profiling, --relaxed-prot-inf, --reannotate, --peak-center, -- no-ifs-removal`. Raw precursor intensities from DIA-NN were then normalized using calibration peptides spiked into each sample prior to MS injection and aggregated to protein intensities using the MaxLFQ algorithm in the iq R package^31^, keeping precursors from different nanoparticles separate. Plate-level batch effects were corrected using the removeBatchEffect function from the limma R package^32^. Further preprocessing includes log2-transformation and missing value imputation using a sequential k-nearest neighbors (KNN) algorithm^33^. While MS-Seer provides additional peptide-level information for each protein, it was not used in this study to enable direct comparison between three platforms.

### Limit of detection and missingness

Limit of detection (LoD) is defined as the lowest protein concentration likely to be reliably distinguished from analytical noise (i.e., the signal produced in the absence of protein)^34^. For LoD calculations, we adopted the formulae recommended by each platform, using readouts from blank samples. For SomaScan data, the LoD was calculated as Median (Blank) + 4.9 * MAD (Blank). For Olink data, the LoD was derived as Median (Blank) + 3 * SD (Blank) or Median (Blank) + 0.2, whichever value is highest, using function called olink_lod function from OlinkAnalyze R package. The proportion of data below LoD is termed as missingness.

To enable fair comparison between platforms, we also applied consistent formulae based on the Clinical and Laboratory Standards Institute (CLSI) definitions^34^ to compare the overall proportion of measurements below the LoD for each platform. These formulae included: Formula 1: LoD = Mean (Blank) + 3.3 * SD (Blank); and Formula 2: LoD = Mean (Blank) + 1.65 * SD (Blank) + 1.65 * SD (Lowest 5% samples).

For the MS data, proteins were identified and quantified only if they met an FDR threshold of less than 1%; otherwise, they were recorded as NA. In our analysis, we considered missing values as equivalent to measurements below the LoD.

For sample-level missingness, we estimated the proportion of analytes with measurements below LoD for each sample, and reported the distribution and median missingness across all samples. For protein-level missingness, we estimated the proportion of samples with measurements below LoD for each analyte, and reported the numbers of analytes with missingness at or beyond specific levels (= 0%, > 20%, > 50% and = 100%). The primary analyses were conducted using all samples (both the first technical replicates and single samples [those without replicates]) and all analytes available for each platform. For LoD-based stratified analyses, we categorized the analytes being measured in each platform as Above LoD or Below LoD based on a 20% threshold, i.e., if an analyte was below LoD (or missing) in more than 20% samples, it was classified as Below LoD and vice-versa.

### Technical reproducibility

For SomaScan11K and OlinkHT, to assess technical reproducibility at the sample level, for each sample with triplicates, we calculate the mean pairwise Pearson correlation among technical replicates. At the protein level, each protein is measured in triplicate across 100 samples, producing three 100-by-1 measurement vectors. We then compute the average pairwise Pearson correlations among these vectors to obtain the protein-level mean Pearson correlation. The above method also applies to MS-Seer data, but only 50 samples have duplicates in this case.

### Coefficient of variation (CV)

We estimated the coefficients of variation (CVs) for technical/longitudinal replicates on the same plate (intra-) and different plates (inter-) for the affinity-based platforms (SomaScan11K and OlinkHT). The CVs for MS-Seer platform were calculated based on technical replicates irrespective of intra/inter-plate runs. The formula used for CV calculation for each platform are as follows:

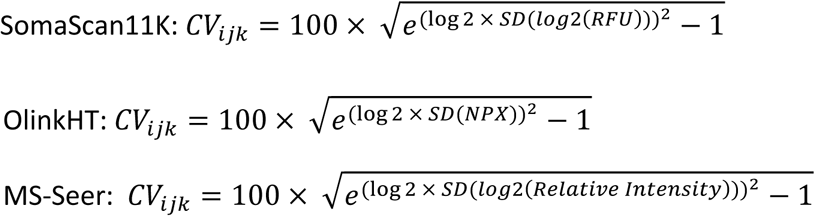

The CVs for each analyte were then averaged across all samples (with technical/longitudinal replicates) to obtain a single CV for each analyte being measured. We then looked at the distribution of these CVs for each platform and reported the CV quantiles. The CVs were calculated for all samples and all analytes available for each platform and also for only the common samples and common analytes shared by all the three platforms.

### Correlation between platforms

We compared two proteomic platforms by examining protein-wise correlations for a set of shared proteins measured in common samples. Proteins were aligned across platforms based on their provided UniProt identifiers, ensuring that only columns corresponding to the same protein in both datasets were compared. Spearman’s rank correlation coefficient was then calculated for each matched protein pair, providing a measure of the agreement between the platforms. The distribution of this calculated correlation for all common proteins between the platforms gives us the observed correlation distribution. To establish a background distribution of correlation values, we performed a permutation-based null experiment in which the protein labels for one platform were shuffled randomly, and Spearman correlations were recalculated for each protein under this randomized mapping. This procedure was repeated 1,000 times, giving us a distribution of correlation values expected under random pairing. The mean of these permuted correlations serves as the background correlation estimate. Observed correlations surpassing the 99th percentile of the background distribution was considered indicative of a strong association between the two platforms.

### Phenotypic association analysis

We evaluated the association of proteins with three key phenotypes: age, sex, and BMI, using generalized linear regression models, adjusting for ethnicity and the two other phenotypes not being tested. Age and BMI were rank inverse-normal transformed prior to association tests. We also evaluated the number of phenotypic associations stratified by above/below LoD proteins and assay sets as described previously (for SomaScan11K and OlinkHT). This analysis was performed across all proteins and common samples, as well as common proteins and common samples, for the two affinity-based platforms (SomaScan11K and OlinkHT). When comparing the associations for ANML data with pre-ANML data, we calculated the associations using all 500 samples and 10,675 aptamers.

### Genotypic association analysis

We performed GWAS using 30X whole-genome sequencing data^35^, filtering variants for missingness <2%, minor allele frequency (MAF) >0.05, and Hardy-Weinberg equilibrium p-value >1×10⁻⁶. Association testing was conducted on 197 samples using the linear mixed model implemented in GCTA-fastGWA^36^, adjusting for age, sex, and the top 20 genetic principal components. Protein expression levels were rank-based inverse normal transformed prior to analysis.

To identify independent association signals, we applied LD-based clumping to GWAS summary statistics using PLINK^37^ with a p-value threshold = 5×10^-8^, LD-R² threshold of 0.01 within a 1Mb window around index variants. Independent genetic variants were classified as *cis* if they were located within ±1 Mb of the gene body encoding the corresponding protein, and *trans* otherwise. A genome-wide significance threshold of p < 5×10⁻⁸ was used for identifying *cis* associations. For *trans* associations, Bonferroni correction was applied based on the number of proteins tested: p < 5.2×10⁻¹² for SomaScan11K (9,563 UniProt IDs) and p < 9.2×10⁻¹² for OlinkHT (5,401 UniProt IDs). To control for multiple analytes targeting the same protein, we also report the number of unique proteins with at least one significant association, i.e. one analyte per UniProt ID. For this, we selected the analyte with the maximum number of associations or a random analyte in case of ties. We also evaluated the number of genotypic associations stratified by above/below LoD proteins and assay sets as described previously (for SomaScan11K and OlinkHT). To allow for a fair comparison, we also analyzed the number of associations for only common samples and common proteins across SomaScan11K and OlinkHT. When comparing the ANML data with pre-ANML data, we calculated the number of genotypic associations using all 500 samples and 10,675 aptamers.

### Fine-mapping of *cis*-pQTLs

We performed statistical fine-mapping using **SuSiE-X**^38^ (single population) to identify putative causal associations. For each lead protein-SNP pair, we extracted summary statistics for all variants within a 1.5Mb window centered on the sentinel variant. Fine-mapping was performed for each locus, with the credible set coverage level set to 90% and p-value threshold of 5e-08. Linkage disequilibrium (LD) matrices were computed from the same genotype data used for the GWAS.

Credible sets were reported if the posterior inclusion probability (PIP) exceeded the defined threshold (>0.90) and the model converged successfully. Variants within the credible sets with PIP >0.5 were classified as causal variants for the specific protein.

### Concordance of *cis*-pQTL and *cis*-eQTL associations

We queried the cis-pQTLs against the eQTLGen^17^ Phase 1 dataset to identify overlapping signals that were also reported as cis-eQTLs for the same gene. We first merged the datasets based on variant position and gene identifier, and subsequently applied a cis-eQTL significance threshold of p <0.05.

### Spike-in simulation procedure

Twenty quantitative phenotypes with different levels of Spearman correlation with ANML scaling factor in the largest dilution bin (1:5) were selected first, including C-reactive protein (CRP), Insulin, Body mass index (BMI), Triglyceride, Bone Mineral Density (BMD)-Hip, Alanine aminotransferase (ALT), Gamma-glutamyl transpeptidase (GGT), Waist hip rate (WHR), Systolic blood pressure (SBP), Glucose, Total cholesterol, Albumin, Uric acid, BMD-Lumbar, Age, Urea, Creatinine, Na, Cl and High density lipoprotein (HDL). Then for each phenotype, we generated 100 synthetic proteins (50 with positive effect size and 50 with negative) with p-values ranging from 1e-0.5 to 1e-20. To minimize the influence on sample distribution, the chosen proteins from the 1:5 dilution bin to replace have the same direction of effect size (positive or negative). And data for 100 synthetic proteins were replaced in the pre-ANML SomaScan11K data under their original column names while keeping other columns unchanged. Altogether, we developed 2 methods to generate synthetic proteins (details described below) and generated 40 datasets (2 methods × 20 phenotypes), with each file containing 100 synthetic data columns. Finally, SomaLogic helped to run ANML on these new datasets and we checked how ANML influences the effect size beta and p-value in linear regression (phenotype ∼ each simulated protein).

### Simulation method 1

We generated synthetic proteins for each phenotype with desired correlations between the protein RFUs and target phenotypes by reshuffling the sample rows in the original pre-ANML data. First, protein RFU ranks were either perfectly aligned with the phenotype ranks (Spearman ρ = 1) or aligned in the exact opposite direction of the phenotype (Spearman ρ =-1) depending on the desired direction of association. To decrease this correlation and control the p-values for linear regression, a specified fraction (e.g., 10%, 25%, 50%, 80%, 90%) of these ranks were randomly shuffled. To avoid perfect linearity between the remaining unshuffled ranks and the phenotype, and to introduce realistic variability, we added minor, random noise to these unshuffled values. This noise was scaled relative to the standard deviation of the log₂-transformed protein intensities using a defined factor (0.25) so that the magnitude of the noise was strong enough to remove the residual linearity in the unshuffled data. Using the above-described method, we generated 50 proteins with positive and 50 with negative association with each target phenotype, with the desired degree of correlation, while also preserving the distribution pattern of the original protein columns that were replaced.

### Simulation method 2

For each phenotype, inverse normal transformation was applied firstly to derive Z_Phenotype_ which follows a standard normal distribution. Next, a target Pearson correlation ρ was set. Given Z_Phenotype_, we generated Z_Protein_ from a normal distribution with mean ρ Z_Phenotype_ and variance 1-ρ^2^. Then by probability theorem, Z_Protein_ follows a standard normal distribution and has Pearson correlation ρ with Z_Phenotype_. We then mapped Z_Protein_ to the distribution of one original protein in pre-ANML data (by mapping the quantiles) and obtained a simulated protein with correlation close to ρ with Z_Phenotype_. Using this method, we generated simulated proteins for each phenotype while controlling for the linear regression p-value and target Pearson correlation ρ, and preserving the distribution pattern of the original protein columns that were replaced.

### Analysis of ethnic differences

R function *prcomp* was used to perform PCA of proteomic data. Hotelling’s T2 test for multivariate pairwise comparison was used to estimate significant differences in the means of principal components 1 to 5 for each pair of ethnicities. Linear regression adjusting for age and sex was used to determine association of each analyte with ethnicity, and a significance threshold of FDR<0.05 was applied to each pairwise comparison (Indian vs Chinese, Malay vs Chinese, and Malay vs Indian). For the protein P18428 (LBP), we estimated pairwise Spearman correlation with 9 clinical variables BMI, mean systolic and diastolic blood pressure (mean SBP and mean DBP), C-reactive protein (CRP), triglyceride (Trig), total cholesterol (TC), high density lipoprotein (HDL), low density lipoprotein (LDL) and TC to HDL ratio. Correlation analysis was performed for protein measured by OlinkHT, SomaScan11K Pre-ANML, and SomaScan11K ANML, and a p-value threshold of 0.005 (correcting for the 9 tests) was applied to each set of analysis.

### Bioinformatics and statistical analyses

For all analyses, we excluded assays with multiple UniProt IDs, non-human or contaminant proteins, and internal control proteins. Outlier detection was conducted using four criteria: PCA (PC1 & PC2), sample median, sample IQR, and mean Pearson correlation with other samples. Samples deviating more than five standard deviations from the mean in any of these metrics were considered outliers. For technical evaluations involving common proteins across platforms, we retained only one representative analyte (one with the lowest proportion below LoD) for protein whose UniProt ID corresponds to multiple analytes. Pearson correlation was used for within-platform technical metrics whereas Spearman correlation was used for between-platform comparison.

We also stratified the analysis based on the different release generations of the platforms: For SomaScan11K, the SOMAmers were divided into three sets: (set 1) assays in SomaScan 5K, (set 2) assays in SomaScan 7K minus assays in SomaScan 5K, and (set 3) assays in SomaScan 11K minus assays in SomaScan 7K. For OlinkHT, the three sets were: (set 1) assays in Olink Explore 1536, (set 2) assays in Olink Explore 3072 minus assays in Olink Explore 1536, and (set 3) assays in Olink Explore HT minus assays in Olink Explore 3072. The analysis was also stratified by the missingness of proteins, with above-LoD proteins defined as proteins measured in 80% or more samples and below-LoD proteins defined as proteins missing in more than 20% samples.

ANML scaling factors were derived by dividing post-ANML protein RFU by the corresponding pre-ANML protein expression on the original scale, yielding one scale factor for each sample–protein pair. As these factors are highly similar among proteins within a given dilution bin, the median value within each bin was employed as the representative scaling factor for all proteins in that bin.

The relative contributions of all 29 phenotypes to ANML scaling factor prediction were assessed using a random forest–based Boruta feature selection procedure. By iteratively comparing the importance scores of each actual phenotype against those of randomized shadow features, Boruta robustly identified the subset of phenotypes whose importance significantly exceeded that of noise. The confirmed features thus highlight the key traits driving ANML scaling factor estimation.

## Data and code availability

Researchers may apply for access to individual level data through the study Data Access Committee. All summary data have been made available in the supplementary tables. The R and Python scripts for analysis and visualization are available via <u>GitHub</u>.

## Acknowledgements

We thank the participants of the HELIOS study and the HELIOS operation team for recruitment, organization and data/sample collection. We would also like to thank the National Precision Medicine Programme for sharing relevant whole genome sequence data. This study (NTU IRB: 2016-11-030) is supported by Singapore Ministry of Health’s (MOH) National Medical Research Council (NMRC) under its OF-LCG funding scheme (MOH-000271 and MOH-001792) and NMRC through National Cohorts Office (P2022-02-03) and the National Precision Medicine (NPM) Programme. NPM Programme Phase II is supported by the National Research Foundation, Singapore (NRF) under the RIE2020 White Space (MOH-000588 and MOH-001264) and administered by the Singapore Ministry of Health through the National Medical Research Council (NMRC) Office, MOH Holdings Pte Ltd. NPM Programme Phase III is supported by the Singapore Ministry of Health through the NMRC Office, MOH Holdings Pte Ltd under the NMRC RIE2025 NPM Phase III Funding Initiative (MOH-001734). HELIOS is also supported by intramural funding from Nanyang Technological University, Lee Kong Chian School of Medicine and National Healthcare Group.

## Author Contributions Statement

J.C.C and N.S conceived and designed the study. A.B and H.W processed the proteomic data. A.B, H.W, P.R.J, N.S performed data analyses. A.B, H.W, P.R.J, J.C.C, N.S wrote the manuscript. L.D.W, O.B, Z.D, J.F, J.N, J.Lee, E.R, J.Liu, S.MS, E.P, CY.C, B.L, S.S.J, N.B, C.B, M.L, W.Z, X.S, P.T provided critical feedback on data analysis, interpretation of results, and writing of manuscript. All authors reviewed and contributed to the revision of the submitted manuscript.

## Extended Data Figures

**Extended Data Figure 1.**
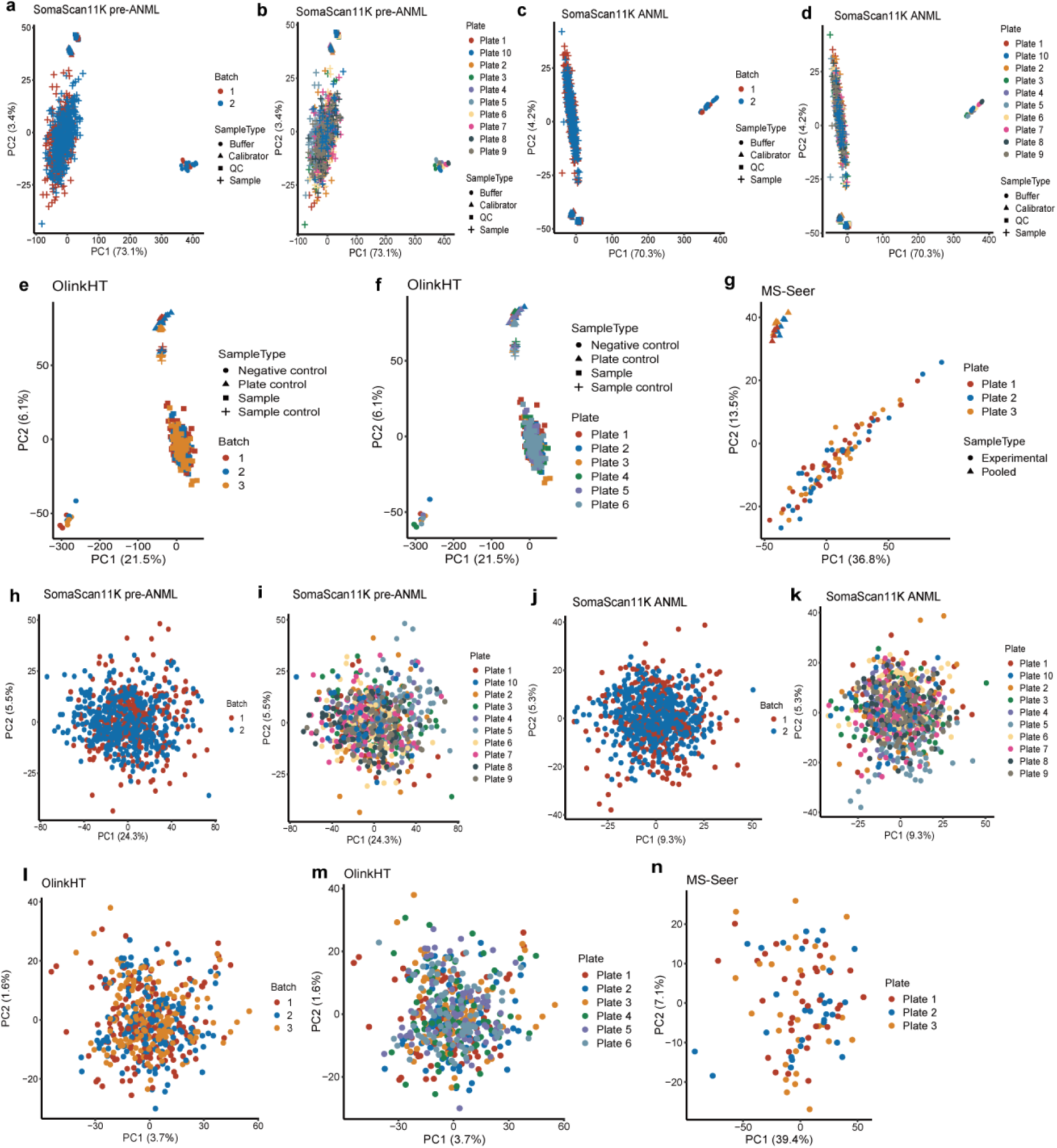
Overview of data across three proteomics platforms. (a-g) PCA plots displaying the overall distribution of experimental and control samples: (a, b) PCA of SomaScan11K pre-ANML data, grouped by Batch (a) and Plate (b); (c, d) PCA of SomaScan11K ANML data, grouped by Batch (c) and Plate (d); (e, f) PCA of OlinkHT data, separated by Batch (e) and Plate (f); (g) PCA of MS-Seer data categorized by Plate. (h-n) Additional PCA plots depict the distribution of experimental samples, labeled by Batch or Plate: (h, i) SomaScan11K pre-ANML (Batch and Plate); (j, k) SomaScan11K ANML (Batch and Plate); (l, m) OlinkHT (Batch and Plate); (n) MS-Seer (Plate).

**Extended Data Figure 2.**
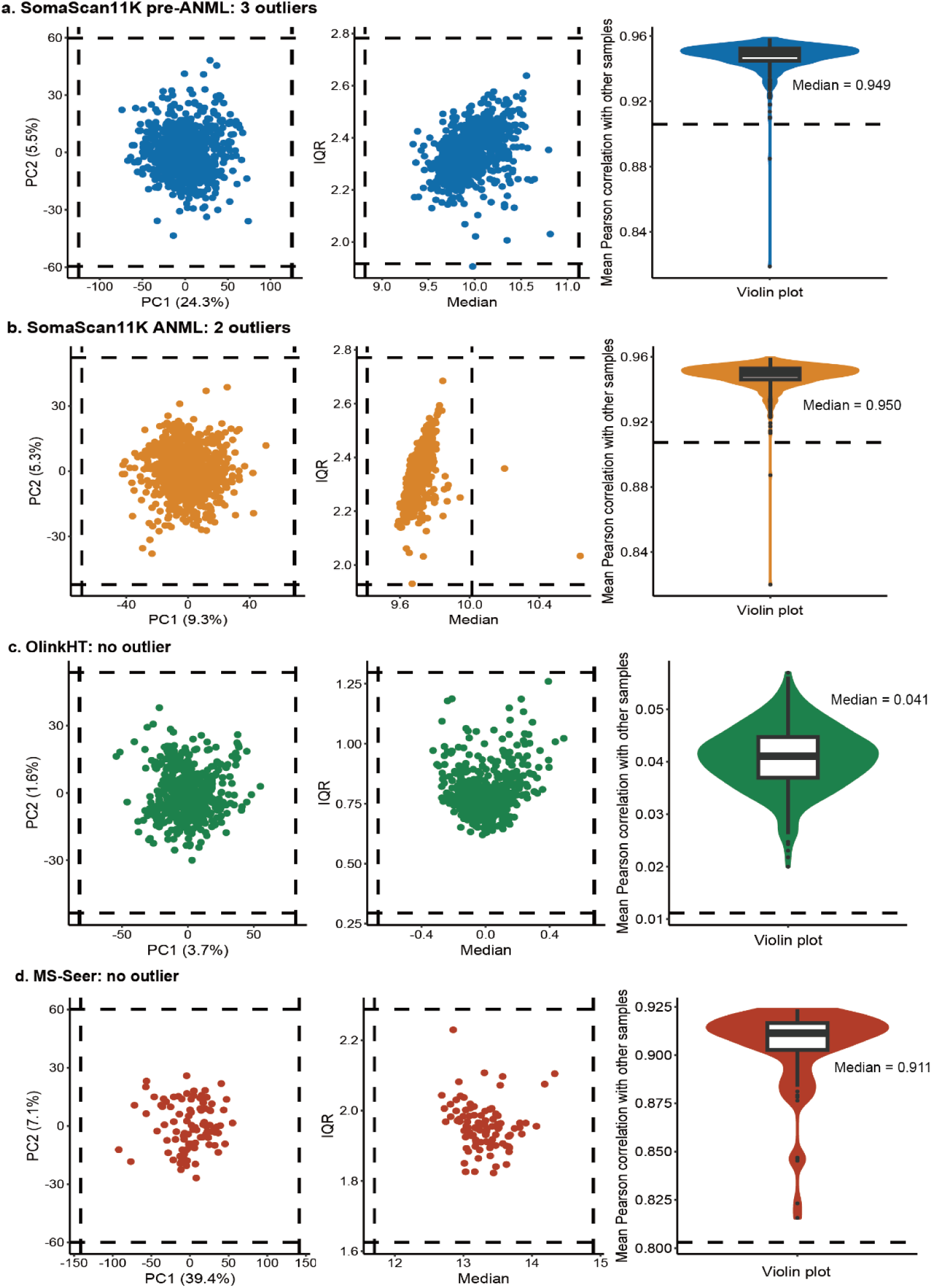
Outlier detection for three platforms. Samples deviating more than five standard deviations from the mean based on four metrics (PC1 & PC2, sample IQR, sample median and mean Pearson correlation with other samples) were considered outliers. The dashed lines in the plot denote the thresholds. (a) Results for SomaScan11K pre-ANML data; (b) Results for SomaScan11K ANML data; (c) Results for OlinkHT data; (d) Results for MS-Seer data.

**Extended Data Figure 3.**
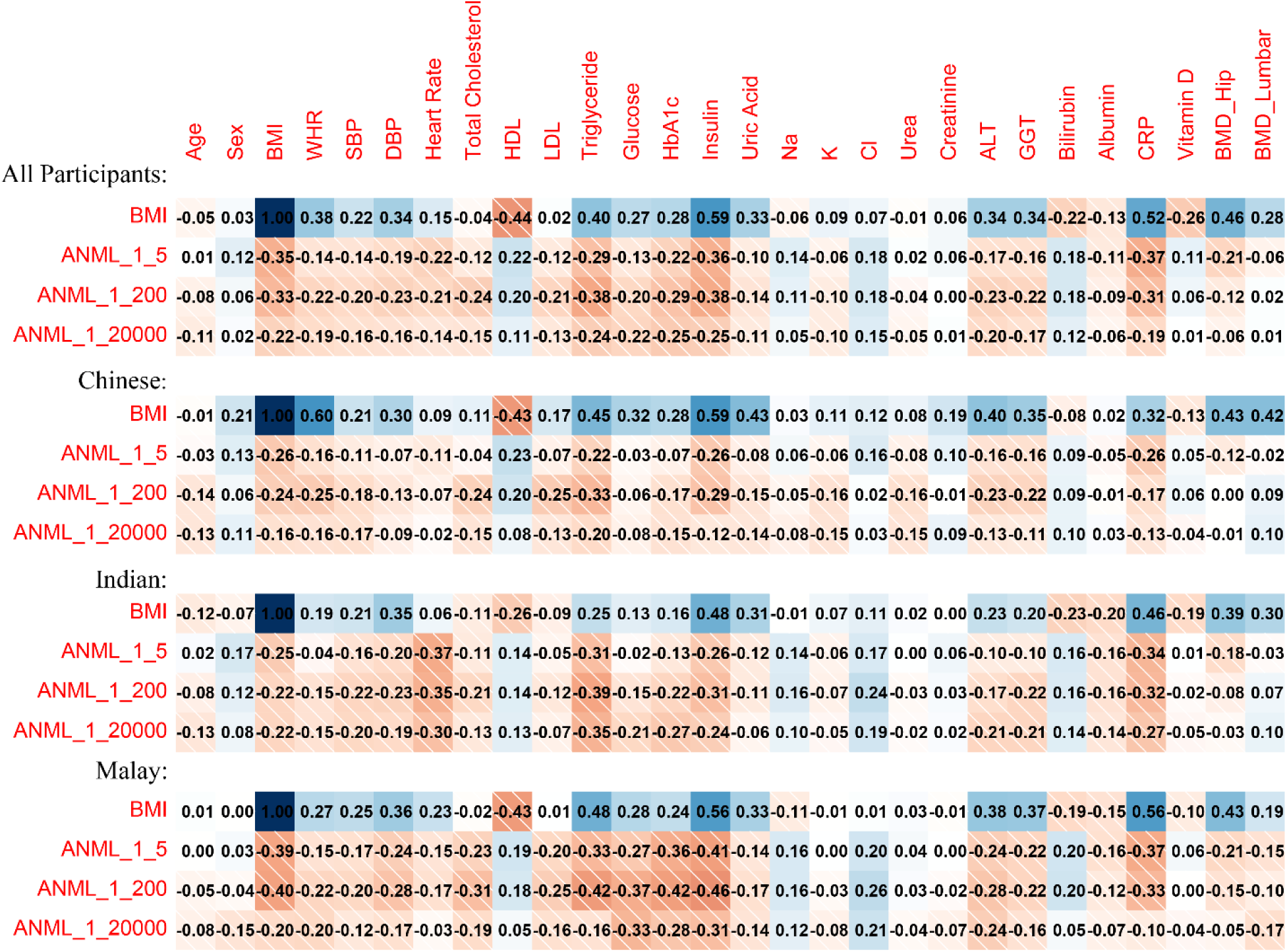
Spearman correlation between ANML scaling factors and a series of phenotypes based on all participants, Chinese, Indian and Malay participants, respectively.

**Extended Data Figure 4.**
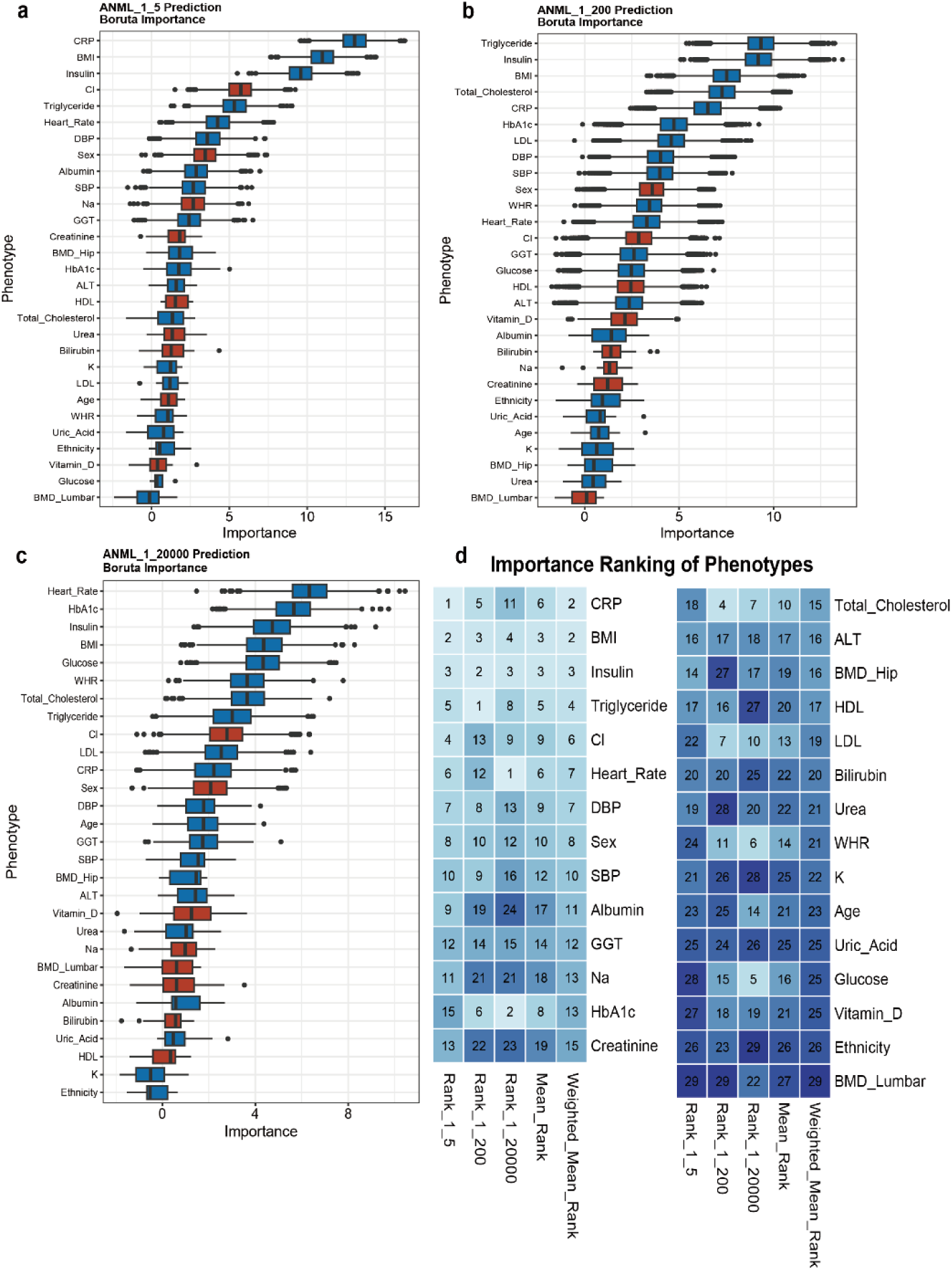
Random forest-based Boruta importance in ANML scaling factor prediction by 29 phenotypes. Phenotype importance in 1:5 dilution bin (a), 1:200 dilution bin (b) and 1:20000 dilution bin (c). Red: Positive association with ANML scaling factor; Blue: Negative association with ANML scaling factor. (d) Summarized importance ranking (according to median importance) across three dilution bins in each prediction. The weighted mean rank is weighted by the numbers of proteins in each dilution bin.

**Extended Data Figure 5.**
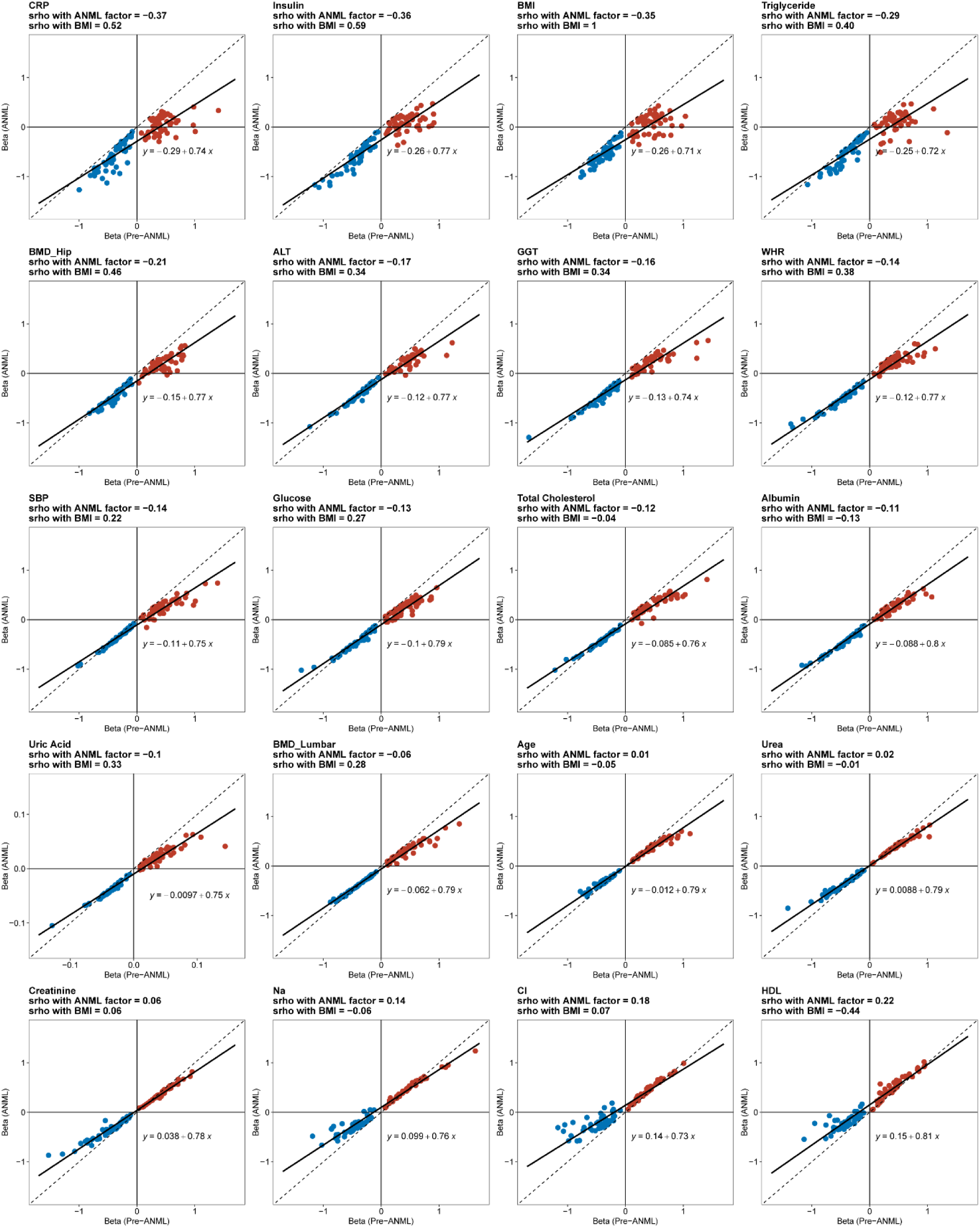
Effect size comparison between pre-ANML and ANML of synthetic proteins across 20 phenotypes (Method 1). The Spearman correlations between each phenotype and ANML scaling factor/BMI are shown in each panel title. The dotted and solid lines represent y = x and fitted line, respectively. Red: pre-defined positive effect size; Blue: pre-defined negative effect size.

**Extended Data Figure 6.**
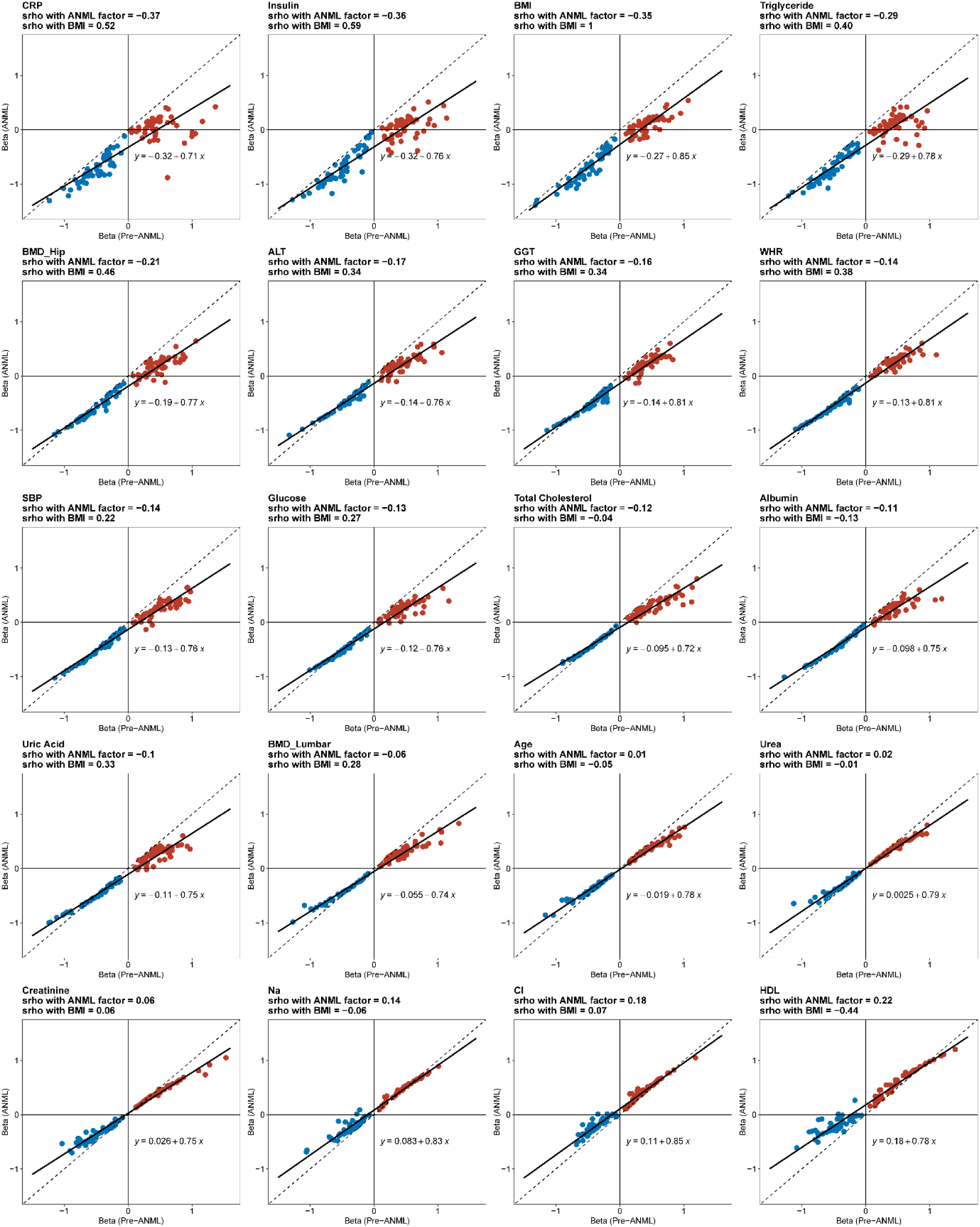
Effect size comparison between pre-ANML and ANML of synthetic proteins across 20 phenotypes (Method 2). The Spearman correlations between each phenotype and ANML scaling factor/BMI are shown in each panel title. The dotted and solid lines represent y = x and fitted line, respectively. Red: pre-defined positive effect size; Blue: pre-defined negative effect size.

